# Phytochemical Screening, Investigation of the toxic and hepatoprotective effect of leaves of *Prunus africana*-using mice model

**DOI:** 10.1101/2024.04.11.588968

**Authors:** Nikodimos Eshetu Dabe, Adane Teshome Kefale, Habtamu Acho Addo, Teshale Worku Dido, Muktar Sano Kedir, Hana Tadesse Afework, Molla Asnake Kebede

## Abstract

**Purpose:** Liver disease is a growing public health problem both in developing and developed countries. Allopathic medicines available for liver disease are not accessible and affordable for 3^rd^ world countries, like Ethiopia. There is a need to search for alternative therapy, in which herbals are widely used in traditional medicine and promising for the development of effective and less costly treatment. As a result, this study aimed to screen phytochemicals and investigate the toxic and hepatoprotective effects of *Prunus africana* which is traditionally used in Benchi sheko and Sheka Zones, Southwest Ethiopia.

**Methods:** Methanolic extracts of *Prunus africana, w*ere used to evaluate the toxicity and hepatoprotective activity in both male and female swiss Albino Mice. Liver injury was induced by carbon tetrachloride (CCl4). The toxicity and hepatoprotective activity were examined biochemically, histologically, and through the analysis of the general features of the study animals. The phytochemical composition was screened qualitatively using standard chemical tests for secondary metabolites. The results were expressed as Mean ± SD, and differences at P < 0.05 were considered significant. Differences between the experimental and control groups were analyzed using one-way analysis of variance (ANOVA) followed by Dunnett’s T-test to determine their level of significance.

**Results:** The extract of *Prunus africana* revealed positive for the presence of flavonoids and Saponins but negative for Anthraquinone glycoside. The current study showed that the median oral lethal dose of the plant was greater than 5000mg/kg. The general behavior of the animal and organ weight, gross morphology, and the biochemical and histological parameters confirmed that the Methanolic leaves extract of this plant is safe during the sub-acute toxicity tests with a dose of 600 and 1800mg/kg. Methanolic extract before and after the CCl4 administration caused a significant reduction in the values of Alanine transaminase, Aspartate transaminase, Alkaline phosphatase, and bilirubin (P<0.05) almost comparable to the standard drug Silymarine. The hepatoprotective activity was supported by histopathological examination of the liver tissue of control and treated animals.

**Conclusion:** The methanolic extract of *Prunus africana* at the test doses did not show significant toxicity. The results of this study also demonstrate that the extract was effective for the prevention of CCl4-induced hepatic damage in Mice. This study thus justifies that, the use of *Prunus africana* in the treatment of liver diseases and points out that this plant warrants further detailed investigation as a promising hepatoprotective agent.

## 1. Introduction

### 1.1. Types, Etiology, Pathophysiology, and Epidemiology of Liver Disease

The liver is an important metabolic and excretory organ. A damaged liver cannot perform all of its functions properly and may not secrete bile acid (1). Chronic liver disease occurs worldwide, regardless of age, gender, region, or race, and remains one of the most serious global health problems, affecting more than 10% of the world’s population (2). In the clinical context, it describes pathological processes in the liver that are associated with progressive destruction and regeneration of the liver parenchyma, which, if left untreated, ultimately lead to liver cirrhosis and hepatocellular carcinoma (3). Liver cirrhosis is the end result of various liver diseases characterized by fibrosis and abnormal liver architecture with regenerative nodule formation, which can lead to various clinical symptoms and complications (4).Cirrhosis was estimated to be responsible for over 1 million deaths worldwide in 2010, accounting for approximately 2% of all deaths worldwide (5). In viral hepatitis in particular, it is estimated that around 250 million people worldwide are affected by hepatitis C and 300 million are carriers of hepatitis B (6).

The signs and symptoms, the natural history, and the rationale of treatment for all forms of liver disease derive from certain relatively simple basic pathophysiologic concepts. All of the basic pathophysiologic mechanisms of both infectious hepatitis and cirrhosis can be classified into two major categories: those that are the result of impairment of hepatocellular and Kupffer cell functions, and those that derive from impairment of hepatic circulation in advanced cirrhosis (1).

Interferon and ribavirin are some of the drugs currently prescribed for viral hepatitis, so far, there is no effective treatment for hepatitis today (2) and the current management of liver disease is not accessible and affordable for many patients residing in the different corners of the globe, particularly those of developing countries, including Ethiopia, leaving traditional medical practice the choice of treatment.

In Ethiopia, there are no conventional drugs available in health facilities for the treatment of liver diseases, hence many affected patients are using herbal preparations to get rid of the disease.

Recently, herbal products have gained attention in treating chronic liver diseases worldwide due to their high abundance, long-lasting curative effects, and few adverse effects (3,4). Herbal drugs play a significant role in the regeneration of liver cells, acceleration of the healing process, and management of many liver disorders (3).

Many folk remedies of plant origin are tested for their potential antioxidant and hepatoprotection against liver damage in experimental animal models. For example, the 70% ethanol extract of leaves of *Peltophorum pterocarpum* at 100mg/kg and 200mg/kg doses significantly reduced the elevated levels of biochemical markers (5). In addition, the extracts of *Adansonia digitata L., Ichnocarpus frutescens (Linn.), Marrubium vulgare, Withania somnifera, Curcuma Longa,* and *Clerodendron Inerme* are some of the few currently explored herbal products for their effectiveness in the management of liver injury secondary to either carbon tetrachloride or acetaminophen (3,6–8). Their effectiveness makes the management safer, cheaper, and affordable for resource-poor settings as they can be cultivated in most segments of the world.

Similarly*, P. africana* is one of the herbs that have traditional claims for the management of hepatic disease in the southwest regions of Ethiopia.

*P. africana*, which belongs to the family Rosaceae, is a widespread evergreen tree, growing at an altitude of 1500–2000 meters, usually 10–25 meters high with alternate leaves, a straight cylindrical trunk, a dense rounded crown, and small white or cream fragrant flowers (Fig 1) (9). In Africa, it is widely used to treat malaria, benign prostatic hyperplasia, stomachache, and fever (10). Some species of this family, *Prunus persica* and *Prunus armeniaca L*. are reported to have hepatoprotective effects (11,12).

**Figure 1:**
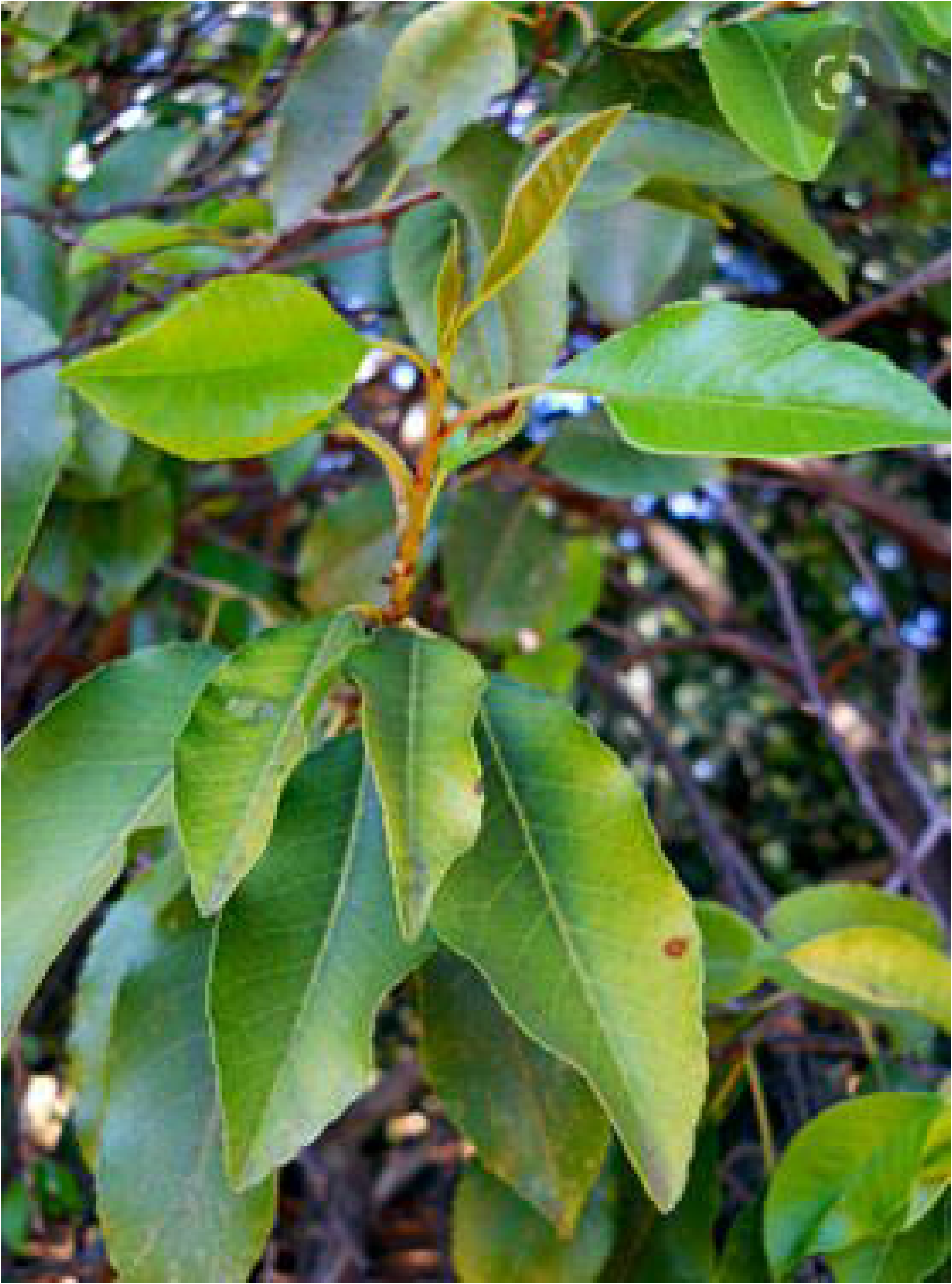
Photographs showing leaves of P. africana taken from Sheka forestry region, Ethiopia, in November 2023.

Despite, the literature trace of its effects on Hepatic disease, it is widely utilized in Bench-Sheko and Sheka zones, Southwest Ethiopia for their traditional claim of the Hepatoprotective effect. Further investigation was needed to strengthen and support the societal practice by scientifically and experimentally proofed evidence so that the world can benefit from these easily available naturally occurring products with better safety profiles.

Therefore, the present study aimed to evaluate the toxic and hepatoprotective effects of *P. africana* traditionally used in the treatment of liver disease in southwest Ethiopia. The result can provide evidence to support the traditional claim of the above plant or to refute the claim. The study can also be used as input for further studies, potentially for the development of new drugs.

## 2. Materials and Methods

### 2.1. Study design

The laboratory-based experiment was employed

### 2.2. Plant Materials and Preparation of Extracts

Fresh plant materials of *P. africana (Hook.f.) Kalkman was* collected based on its ethnobotanical description from forestry regions of Sheka zone, Andracha district, Southwest Ethiopia. The Specimens of the plant were identified by a taxonomist and a few samples were deposited at the National Herbarium of the College of Health Science, Mizan-Tepi University (MTU) with a voucher spacemen number of (105/MTU/PHARM). Fresh parts of the leaves of the plant were cleaned from extraneous materials, dried under shade at room temperature, and grinded by manual crusher and obtained fine particles. Then, It was further processed to obtain the hydro-alcoholic extract as to Ketema M. et al., (13)

The hydro-alcoholic fraction of this medicinal plant qualitatively screened for the presence of secondary metabolites to relate the hepatoprotective activity with the presence or absence of the constituents. Standard chemical tests for secondary metabolites employed (14–16).

**FeCl_3_ test was used to test tannins:** powder of tannins treated with few drops of FeCl_3._ Tannins would give blue-black (hydrolyzable tannins) or green color (condensed tannins).

**To test alkaloids,** the plant extract (0.5gm) would be added to 5ml of dilute H_2_SO_4_ (1%) in a steam bath. The solution would be filtered, and treating the filtrate with the following reagents will produce a corresponding colored precipitate in the presence of alkaloids.

**Mayer’s reagent:** yellowish-white precipitate with potassium mercuric iodide.

**Lead acetate Test was used to test flavonoid:** Extracts were treated with a few drops of lead acetate solution. The presence of flavonoids is shown by the production of a yellow-colored precipitate.

**A Borntrager’s test was used to test anthraquinone:** A little powder was shaken with an immiscible solvent chloroform and then filtered. To the filtrate in a test tube, an equal amount of Ammonium hydroxide is added and shaken. The development of pink, red, or violet colors in the aqueous phase indicates the presence of free anthraquinone.

**A froth test has been used to test for saponins:** Saponins and drugs containing saponins give persistent froth (foam) when shaken with water.

**Libermann Burchard’s test:** The plant powder of extract (0.5gm) dissolved in 2ml chloroform and filtrated. To about 2ml of the filtrate, acetic acid anhydride (2ml) is added. Two milliliters of concentrated H_2_SO_4_ were then added. The development of blue-green color shows the presence of steroids

**Salkowski’s Test:** The extract would be mixed with 2ml of chloroform and filtered. To the filtrate, three drops of concentrated H2SO4 would be then carefully added to form a thin layer. A reddish-brown coloration at the interface would indicate the presence of terpenoids.

**Chemicals used**: 80% methanol, Ferric chloride, sulphuric acid, potassium mercuric iodide, lead acetate, chloroform, Ammonium hydroxide, acetic acid anhydride, normal saline, distilled water, and diethyl ether. All chemicals were purchased from an Importer or company called Fine Chemicals, Addis Ababa, Ethiopia.

### 2.3. Experimental Animals

Swiss albino mice, 2 - 3 months old and weighing 16 - 30gm used in this evaluation. These are procured from the animal house of Mizan Tepi University. They are housed in well-ventilated stainless-steel cages at room temperature (24±2°C) in hygienic conditions under natural light and dark schedules. The relative humidity was 30% to 70% other than during room cleaning, the aim would be 50-60% (8). For feeding, conventional rodent laboratory diets are used with an unlimited supply of drinking water. Then the animals were acclimatized to laboratory conditions for one week before the experiment to alleviate any non-specific stress (8).

### 2.4. Acute and sub-acute Toxicity study

An acute oral toxicity test for the crude 80% methanol extracts was performed according to the Organization for Economic Cooperation and Development guidelines (OECD) (17). As to the guideline, 300mg/kg body weight was selected as the appropriate initial dose for the acute toxicity test and increased to a higher dose of 5000mg/kg body weight. Each of the doses comprised of the hydro-alcoholic leaves extracts. A total of six groups of animals were used (Five experimental and one control) each group consisting of 6 adult female mice. All mice were fasted 3–4 hours before and 1–2 hours after administration of the 80% methanol extracts. Then, mice were observed continuously for 4 hours with a 30-minute interval and then for 14 consecutive days with an interval of 6 hours for the typical indications and symptoms of poisoning. The individual mice were identified by marks on their tails made with different colored permanent ink and through cage markings indicating the group designation and treatment dose. Following, the volume of the extract administered at a time was 1ml/100g body weight of mice. Each animal received a single dose of extract or vehicle in an adjusted final volume. The handling of animals and general parameters was monitored. According to the OECD, the body weight of each mouse was recorded initially, on the 7^th^ day and at the end of the experiment. The differences in the body weight were also recorded.

Then the entire mice were sacrificed with a high dose of anesthetic diethyl ether. All animals in the study were subjected to careful examination of the external surface of the body and abdominal cavities and their contents including the kidneys and liver. Comprehensive gross pathological observations were carried out on these organs to check for any signs of abnormality and the presence of lesions.

The LD_50_ of the extract was determined after recording the deaths of mice. The dose at which 50% of the mice in a group died was considered to be oral LD_50_ (17).

The sub-acute toxicity study was carried out over four weeks (28 days) (18). One-sixth of the maximum tolerated dose of the acute toxicity study (600 mg/kg, body weight) was selected as the lower dose and one-half of this dose was taken as the maximum dose (1800mg/kg) for the sub-acute toxicity test (19). The plant extract is tested on 36 (18 male and 18 female) Mice. There were two experiments (lower dose (600mg/kg) groups and higher dose (1800mg/kg) groups) and one control animal group for each sex. The individual Mice were identified, handled, treated, and analyzed as it is stated in the acute toxicity study section.

The acute and sub-acute administration of the extract were done one after the other to use the results of the acute toxicity study as a baseline for the sub-acute toxicity study.

At the end of the experimental period, animals belonging to each group were weighed on a digital balance and each animal was anesthetized by diethyl ether and blood samples were withdrawn by cardiac puncture. The plain test tubes with no EDTA (Ethylene Diamine Tetra-acetic Acid) were used for biochemical analysis, the blood samples in the plain test tubes were allowed to stand for 3 hours for complete clotting and then centrifuged at 3000 Rpm (Revolution per minute) for 15 minutes using a bench top centrifuge (HUMAX-K, HUMAN-GmbH, Germany). The plasma was withdrawn and transferred into other clean vials. The sera were kept at −20°C until analysis for clinical biochemistry measurements. The concentrations of ALT, AST, total bilirubin, urea, and creatinine were automatically analyzed by (AUTO LAB18 clinical chemistry analyzer, Italy).

Following the collection of blood samples, each animal was sacrificed, the abdominal cavity was opened and the liver and kidney were carefully removed and cleared from any surrounding tissues by normal saline then put on clean paper and weighed quickly on digital balance. The samples from the liver and kidney were preserved in a 10% neutral buffered formalin fixative solution for the histological process.

Stained tissue sections of the liver and kidney were carefully examined under a binocular compound light microscope (LEITZ WETZARE, Germany) and CX41RF, Philippines). Tissue sections from the treated groups were examined for any evidence of histopathological changes concerning those of the controls. After examination, photomicrographs of selected samples of liver and kidney sections from both the treated and control mice were taken under a magnification of x40 objective lens by using (EVOS XL, USA) automated built-in digital photo camera

### 2.5. Hepatoprotective Activity in Experimental Design

Mice were randomly assigned into 5 groups: (one normal control, one positive control, one toxic (induction) control, and 2 test groups) comprising six animals per group for the hepatoprotective activity test. Normal controls were treated with the vehicles/distilled water, 1ml/100gm orally for 15 days. Positive controls were administered with the standard drug, Silymarin 100 mg/kg for 15 days, and Induction control was administered with CCl4 (50% CCl4 dissolved in liquid paraffin in 1:1) 2 ml/kg IP(intraperitoneal) on the 4^th^, 9^th^ and 13^th^ day of the experiment. Among the test groups (groups 4 and 5) were administered with 300 mg/kg and 600 mg/kg doses of 80% methanol leaf extracts daily for 15 days respectively. Both the test groups were given CCl4 (2 ml/kg) on the 4^th^, 9^th^, and 13^th^ day via IP injection. Doses were determined using data from the acute and Sub-Acute toxicity tests. The route of administration for the test sample, standard drug, and vehicles was orally by using oral gavage and the volume administered was 1ml/100gm body weight. On the 16^th^ day, the mice were sacrificed under light ether anesthesia. Each mouse had a heart puncture to obtain a blood sample, which was then collected in sterile, heparinized centrifuge tubes. The tubes were then separated at 3000 Rpm for 15 minutes in a centrifuge. After the liver function tests were completed, the levels of ALT (Alanine Aminotransferase), AST (Aspartate Amino transferase), ALP (Alkaline Phosphatase), and total bilirubin were measured in clear serum by using the automatic analyzer and commercial assay kits. In addition, the liver of each Mouse was reaped, weighed, and examined for gross and microscopic pathology.

The percentage protection of the extracts’ hepatoprotective activity can be computed as follows: % protection = (a-b)/(a-c)*100 where ‘a’ represents the mean marker value produced by the hepatotoxic; ‘b’ represents the mean marker value produced by the toxin plus test material; and ‘c’ represents the mean marker value produced by the vehicle control (16).

### 2.6. Data processing and analysis

All the data was packed and analyzed by SPSS version 22 statistical software. All values of parameters were expressed using appropriate statistical summary measures (frequency, median, mean ± SD). Results over time would be compared between control and treated groups by analysis of covariance. The results were expressed as Mean ± SDE, and differences at P < 0.05 were considered significant. Differences between the experimental and control groups were analyzed using a one-way analysis of variance (ANOVA), followed by Dunnett’s T-test to determine their level of significance.

## 3. Result

### 3.1. Yield of plant sample extraction

Leaves of *P. africana* (Hook.f.) Kalkman was shaded, and dried at room temperature. Then, they were crashed into coarse powders. A total of 800gm was weighed by electronic balance and macerated by using 80% methanol. The macerated chamber was placed at room temperature for 72 hours with occasional stirring. After 72 hours the extract was first filtered by using muslin cloth then the filtrate was further filtered by using Whatman number 1 filter paper. Then it was kept at 70^0^c over a water bath until the menstrum evaporated. It was macerated again twice. Finally, we obtained 195.6 grams of extract. Then, the dried extract was kept in the refrigerator until used for the experiment.

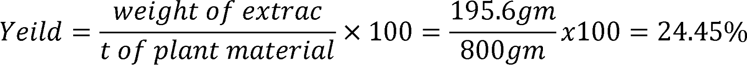

### 3.2. Qualitative chemical tests for secondary metabolites of methanolic leaf extracts of *P. africana*

In this research, the presence or absence of different types of secondary metabolites in hydro-alcoholic extract in *P. africana* was checked through various preliminary qualitative phytochemical tests. Accordingly, the crude extract obtained from *P. africana* using 80% methanol showed positive and negative results (**Table 1**).

**Table 1:** Qualitative chemical tests for the presence/absence of various phytoconstituents of the crude methanolic extracts of *P. africana*.

| Constituents | Test | *P. africana* |
| --- | --- | --- |
| Alkaloids | Mayer's | Positive |
| Flavonoids | Lead Acetate | Positive |
| Anthraquinone glycoside | Borntagor's | Negative |
|  | Modified Borntagor's | Negative |
| Saponins | Foam test | Positive |
| Steroids | Salkowski's | Negative |
| Tannins | Lead Acetate | Positive |
| Terpenoids | Salkowski's | Positive |

### 3.3. Acute oral toxicity tests

Following the third day of administering the 5000 mg/kg dose of *P. africana* methanolic extract, one death was reported. In addition, once the extract was given to the remaining mice at a dose of 5000 mg/kg, symptoms such salivation, diarrhea, lethargy, and irritation were noted. In contrast, these symptoms steadily decreased and disappeared entirely throughout the first week of treatment. This suggested that this plant’s oral LD50 may exceed 5000 mg/kg dosage. The morphological traits of the fur, eyes, and nose seemed normal. There were no signs of tremors, convulsions, salivation, diarrhea, lethargy, or strange behaviors like walking backwards or self-mutilation. The Mice had normal gait and posture, response to sensory stimuli, and grip strength. Except for the final week of extract administration at a dose of 5000 mg/kg, there was no discernible difference in body weight between the treatment and control groups (Table 2). Daily variations were seen in the food and water consumption of the control group of animals.

**Table 2:** Mean body weight of Mice administered with *P. africana* Hook f. extract as compared to the controls during two weeks of observation (expressed as mean ± SDE, n = 6)

| Group | Dose(mg/kg) | Initial mean body weight (gm) | Mean Body weight at the end of week 1 (gm) | Mean Body weight on day 14 (gm) |
| --- | --- | --- | --- | --- |
| I | 300 | 23.4±2.3 | 27.34 ± 2.9(0.54) | 31±2.8(0.56) |
| II | 600 | 21.6±2.13 | 23.5 ± 2.4(0.52) | 25.3±2.1(0.68) |
| III | 1200 | 20.8±1.9 | 22.1 ± 1.2(0.9) | 25.4 ± 1.9(0.5) |
| IV | 3600 | 21.9±2.4 | 23.47 ± 2.3(0.4) | 26.65±2.35(0.39) |
| V | 5000 | 23.2±1.7 | 20.1 ± 2.22(0.04)* | 25.22±2.87(0.48) |
| VI | Control(normal saline) | 22.5±2.8 | 25.85 ± 2.43 | 29.5 ± 2.6 |

The figures under brackets indicate p-values, n =number of mice per group, \* = $p \leq 0.05$

### 3.4. Sub-acute toxicity tests

#### 3.4.1. Effects of sub-acute administration of the extracts on behavior, gross pathology, and body weight

Mice that received repeated oral dosages of *P. africana* extract at 600 mg/kg and 1800 mg/kg body weight did not exhibit any behavioral changes in comparison to the control group during the course of a 28-day sub-acute toxicity test. No aberrant gross findings were observed in any of the experimental or control groups regarding the skin, eyes, liver, or kidneys. Furthermore, during the course of the trial, no deaths were caused by poisoning.

Table 3 demonstrates that almost all male mouse groups experienced a progressive increase in body weight over the course of the investigation. The pattern of body weight gain in each of the experimental and control mouse groups showed no discernible variation.

**Table 3:** Mean body weights (in grams) of male Mice administered with 600 and 1800mg/kg *P. africana* extract during the consecutive four weeks of observation as compared to the controls.

| Effect of *P. africana* extract on body weight of various groups of male Mice (expressed as mean ± SDE, n = 6) | | | |
| --- | --- | --- | --- |
| Week | Control | 600mg/kg | 1800mg/kg |
| Initial mean body weight | 21.5±1.6 | 22.7±2.7 | 21.6±1.6 |
| 1 | 23.7±1.3 | 24.1± 1.6(0.35) | 23.3±1.4(0.79) |
| 2 | 25.2±1.7 | 27.7±1.8(0.16) | 26.3±1.1(0.80) |
| 3 | 28±1.3 | 29.6± 1.2(0.39) | 28.1±1.5(0.74) |
| 4 | 32±2 | 33.2±1.8(0.16) | 32.3±2.2(0.50) |

*The figures under brackets indicate p-values, n – number of mice per group*

The changes in the mean values of the body weights of female Mice administered with 600mg/kg and 1800mg/kg of 80% Methanolic extract of *P. africana* as compared to the controls during the four weeks of the study period is displayed in **Table 4**. The change in body weight was not significantly different in both cases as compared to that of the control.

**Table 4:** Mean body weights (in grams) of female Mice administered with 600 and 1800mg/kg *P. africana* extract in consecutive four weeks of observation as compared to the control.

| Effect of *P. africana* extract on body weight of various groups of female Mice (expressed as mean ± SDE, n = 6) | | | |
| --- | --- | --- | --- |
| Week | Control | 600mg/kg | 1800mg/kg |
| Initial mean body weight | 18.2±1.3 | 19.46±2.1 | 18.73±1.8 |
| 1 | 20.5±2 | 21.89 ± 1.2 (0.52) | 21.3 ± 2(0.54) |
| 2 | 22.2±1.7 | 24.4±1.7(0.6) | 23.8 ± 1.5(0.5) |
| 3 | 23.9±1.87 | 27.8± 1.6 (0.9) | 27.3± 1.7 (0.8) |
| 4 | 28.1±1.54 | 30.8±1.1(0.87) | 29.9±1.64(0.6) |

*The figures under brackets indicate p-value, n – number of mice per group*

#### 3.4.2. Effect of sub-acute oral administration of *P. africana* on the gross morphology of liver and kidney of Mice

The liver and kidney of the male and female experimental and control mice were removed at the end of the study, weighed, and visually examined for any histological alterations in the tissue architecture and color. Both the control and experimental mice’s livers and kidneys revealed normal architecture, no color changes, and no morphological abnormalities upon visual gross examination.

The weight of the liver and kidneys did not significantly change after treatment with *P. africana* extract, as shown in Table 5, as compared to mice given distilled water at doses of 600 mg/kg and 1800 mg/kg.

**Table 5:**
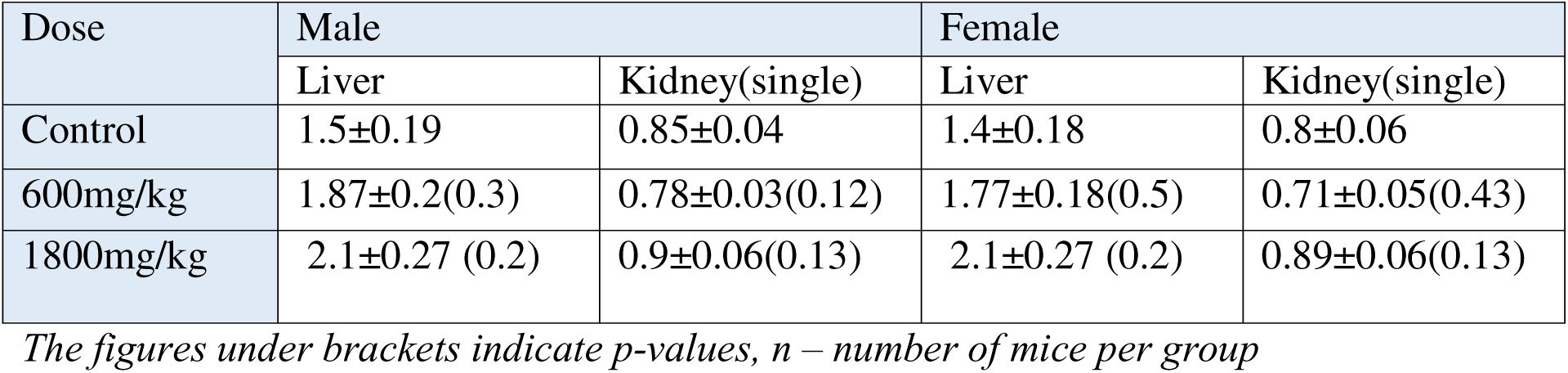
Mean liver and kidney weight of male and Female Mice (in gram; mean ± SDE) Sub-Acutely dosed with *P. africana* as compared to the controls.

| Dose | Male |  | Female |  |
| --- | --- | --- | --- | --- |
|  | Liver | Kidney(single) | Liver | Kidney(single) |
| Control | 1.5±0.19 | 0.85±0.04 | 1.4±0.18 | 0.8±0.06 |
| 600mg/kg | 1.87±0.2(0.3) | 0.78±0.03(0.12) | 1.77±0.18(0.5) | 0.71±0.05(0.43) |
| 1800mg/kg | 2.1±0.27 (0.2) | 0.9±0.06(0.13) | 2.1±0.27 (0.2) | 0.89±0.06(0.13) |

*The figures under brackets indicate p-values, n – number of mice per group*

#### 3.4.3. The Sub-acute Toxic Effects of Methanolic leaf extract of *P. africana* on Biochemical parameters

The Methanolic leaf extract of *P. africana* did not produce a significant change in any of the biochemical parameters: ALT, AST, bilirubin, creatinine, and urea levels in both male and female groups at doses of 600mg/kg and 1800mg/kg when compared with the control group.

However, following the administration of the methanolic extract of *P. africana,* the serum levels of ALT and AST were non-significantly increased in female treated mice at a dose of 1800mg/kg by 25% and 23 % respectively. In addition, the total bilirubin level also non-significantly increased at both 600mg/kg and 1800mg/kg doses by 17% and 25% respectively. In the Renal function test, urea also non-significantly decreased by 6.8% at 1800mg/kg dose of the female group (**Table 6**).

**Table 6:** Effect of 600 & 1800mg/kg Methanolic leaf extract of *P. africana* on biochemical parameters in both male and female Mice as compared to the control group (expressed in mean ± SDE, n = 6)

|  | Parameters | Control | 600mg/kg | 1800mg/kg |
| --- | --- | --- | --- | --- |
| Male Groups | AST(IU/L) | 119.75±30.8 | 147±1.45(0.78) | 151±13.7 (0.16) |
| | ALT (IU/L) | 71.25±26.1 | 101.3±2.07(0.84) | 116 ± 34.9 (0.17) |
| | Total Bilirubin | 0.79±0.09 | 0.81±0.08(0.47) | 0.8 ± 0.13 (0.32) |
| | Urea (mg/dL) | 47.75±5.4 | 48±7(0.843) | 53± 3.2(0.70) |
| | Creatinine (mg/dL) | 0.98±0.08 | 1.04±0.02(0.46) | 1.01± 0.13 (0.55) |
| Female Groups | AST(IU/L) | 101.6±48 | 136±33(0.92) | 146±12.7(0.12) |
| | ALT (IU/L) | 63.3±57 | 83±23.5(0.84) | 118±13.8(0.26) |
| | Total Bilirubin | 1.14±0.36 | 1.84±0.03(0.75) | 2.24±15(0.65) |
| | Urea (mg/dL) | 44.0±8.6 | 41.18±13(0.68) | 49.8±4.5(0.92) |
| | Creatinine (mg/dL) | 0.80±0.2 | 0.88±0.14(0.61) | 1.02±1.12(0.78) |

The figures under brackets indicate p-values, n – number of mice per group

#### 3.4.4. Effects of Methanolic leaf extract of *P. africana* on the liver tissue

Microscopic examination of liver sections from control mice showed the normal architecture of structural units of the hepatic lobules, formed by cords of hepatocytes separated by hepatic sinusoids (Fig 2A and 2B). The central vein and portal area containing branches of the hepatic artery, bile duct, and portal veins were maintained with their normal appearance. In comparison to the control, the general microscopic architecture of sections of liver tissue from the mice treated with the extracts at 600mg/kg dose appeared to be not significantly affected after the 28 days of administration (Fig 2C). However, a liver section of mice treated with the plant *P. africana* with a dose of 1800mg/kg body weight/day showed pyknotic nucleus in hepatocytes and in some areas of the liver sections of mice showed perivascular leukocytic cellular infiltration in the central and portal area (Fig 2D). Generally, there is no histopathological difference between male and female mice of the liver section related to the treated dose.

**Figure 2:**
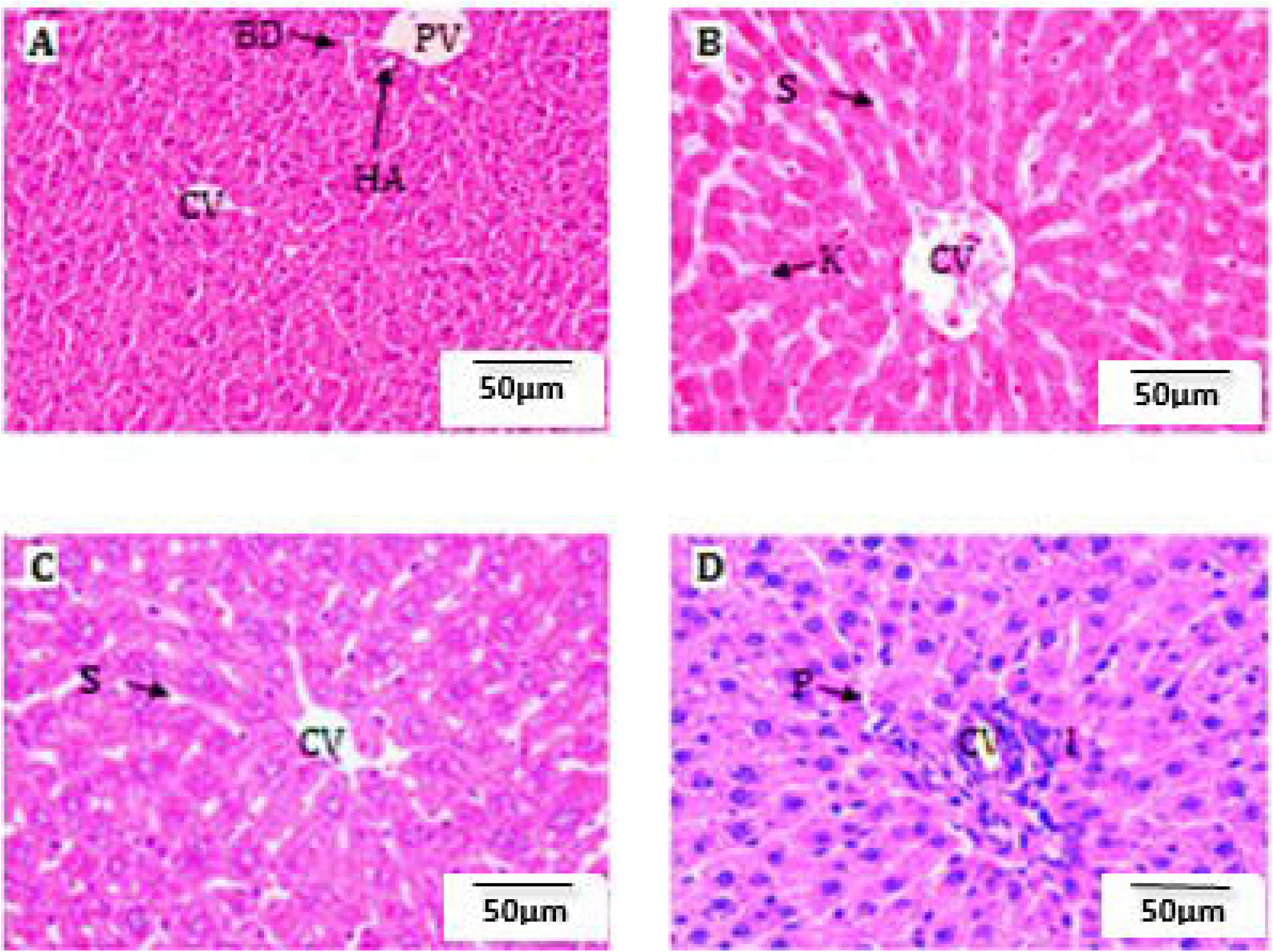
Photomicrographs of liver sections of control Mice (A&B), Mice administered with the extract of 600 (C) & 1800 mg/kg (D) of P. africana extract, Sections are from female Mice. CV = Central vein, E = Endothelial cells, PV = Portal vein, BD = Bile duct, HA = Hepatic artery, K=Kupffer cells, I= leukocytic Infiltration, S= sinusoids, P=Pyknosis. (Sections were stained with H&E, X400).

#### 3.4.5. Effects of Methanolic leaf extract of *P. africana* on histology of the kidney

Histopathological examination of kidney sections of mice treated with the methanolic leaves extracts of *P. africana* at both 600mg/kg (Fig 3C) and 1800mg/kg doses (Fig 3D) indicated no structural disturbance as compared to the control mice (Fig 3A & 3B). The microscopic architecture of the kidneys in treated mice had a similar appearance to that of the controls in which renal corpuscles maintained their normal size of urinary space and normal tubular structures were observed with no sign of congestion. However, mononuclear lymphocytic infiltrations were observed in sections of the kidneys of the mice treated with *P. africana* with a dose of 1800 mg/kg body weight (Fig 3D). There is no histopathological difference between sections of the kidneys of male and female mice related to the treated dose.

**Figure 3:**
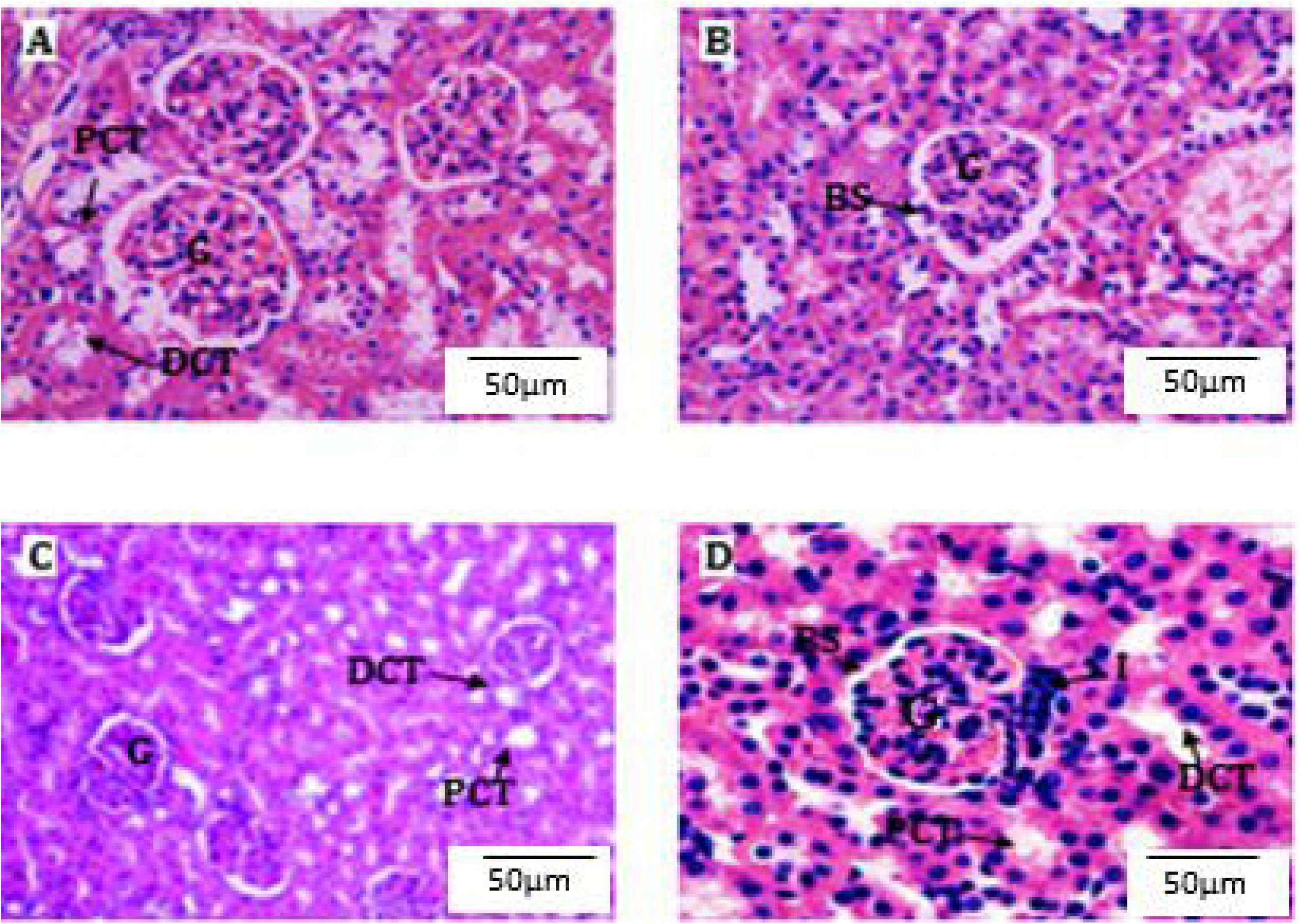
Photomicrographs of Kidney sections of control Mice (A&B), Mice administered with 600mg/kg (C) & 1800mg/kg (D) extract of P. africana, Sections are from female Mice. G = Glomerulus, BS = Bowman Space, DCT = Distal convoluted tubule, PCT = proximal convoluted tubule, I = leukocytic Infiltration. (Sections were stained with H&E, X400).

### 3.5. Hepatoprotective Activity

#### 3.5.1. Effect of methanolic extract of *P. africana* on Body Weight, Absolute & Relative Liver Weight, and Serum Biochemical Markers of Liver

The change in body weight during the two-week test of the Hepatoprotective effect of *P. africana* is shown in **Table 7**. The CCl4-treated control group had a reduced mean body weight (p < 0.05) as compared with those of the normal control group. There was a non-significant weight gain in normal, standard control, and treatment groups at doses 300 & 600mg/kg.

**Table 7:** Effects of methanolic extract of *P. africana* leaves on different liver-specific variables in control and experimental groups of animals. (Expressed in mean ± SDE, n = 6)

| Treatment groups and liver-specific variables | Normal control(1ml/kg normal saline) | Induction control(2ml/kg 50% of CCL4) | Silymarin 100mg/kg ip + 2ml/kg CCL4 | 300mg/kg P. africana + 2ml/kg CCL4 | 600mg/kg p. africana + 2ml/kg CCL4 |
| --- | --- | --- | --- | --- | --- |
| Initial mean body weight | 21 | 20.98 | 20.31 | 19.39 | 21.61 |
| Final mean body weight | 25.7±0.4 | 17.97±0.34<sup>d</sup> | 22.64±0.23<sup>f</sup> | 21.53±0.24<sup>f</sup> | 24.8±0.31<sup>f</sup> |
| Liver weight(g) | 2.43 ± 0.17 | 2.6 ± 0.3 | 2.28 ± 0.13 | 2.11 ± 0.11 | 2.43 ± 0.09 |
| Relative liver weight (%) | 9.4±0.9 | 12.2±1.3<sup>ad</sup> | 10±1.4<sup>f</sup> | 9.8±0.8<sup>f</sup> | 9.7±0.92<sup>f</sup> |
| ALT(IU/L) | 63.4 ± 2.01 | 387 ± 17.2<sup>a</sup> | 69.47 ± 3.2<sup>f</sup> | 174.4 ± 3.8<sup>dc</sup> | 66.5 ± 2.74<sup>c</sup> |
| AST(IU/L) | 69.3 ± 2.49 | 342±15.87<sup>a</sup> | 82.54±8.9<sup>f</sup> | 154.4±14.7<sup>d</sup> | 78.27±12.4<sup>c</sup> |
| ALP(IU/L) | 147.02 ± 1.92 | 276.3±14<sup>a</sup> | 153±9.7<sup>f</sup> | 189.82±15<sup>dc</sup> | 158±10.3<sup>c</sup> |
| Bilirubin (mg/dl) | 0.56 ± 0.02 | 1.82±0.12<sup>a</sup> | 0.65±0.04<sup>f</sup> | 0.83±0.08<sup>dc</sup> | 0.61±0.07<sup>c</sup> |

<sup>a</sup> Compared with normal control P< 0.001. <sup>b</sup> Compared with standard P< 0.001. <sup>c</sup> Compared with induction P< 0.001. <sup>d</sup> Compared with standard P< 0.05. <sup>e</sup> Compared with induction P< 0.01. <sup>f</sup> compared with induction & normal control P>0.05

Effects of *P. africana* on different liver-specific variables in control and experimental groups of animals are also shown in **Table 7**. The CCl4 treated groups significantly (p < 0.05) increased in their relative liver weight as compared to the normal control and standard control.

The biochemical results observed in the treatment of *P. africana* concerning induction of hepatotoxicity using CCl4 were also given in Table 7. Mice treated with CCl4 developed significant (P< 0.05) liver damage and it was well indicated by elevated levels of hepatospecific enzymes like ALT, AST, and ALP in serum. A marked elevation in total bilirubin level (P< 0.05) was observed in the group treated with CCl4 when compared with the normal values and the control groups.

The groups that received the treatment of the methanolic extraction of *P. africana* at dose levels of 300 and 600mg/kg body weight significantly controlled the change in the biochemical parameters. The extract at a dose level of 600mg/kg exhibited a sharp decrease in the serum enzyme levels and the effect was comparable with the standard group treated with silymarin (100mg/kg).

In comparison to the lower dosages (300 mg/kg), the higher dose (600 mg/kg) showed higher hepatoprotective capability. After 600 mg/kg of *P. africana* methanolic extract, the following liver chemistry biomarkers show lowered levels: ALT by 99%, AST by 96.6%, and ALP by 91.5%. ALT and AST were reduced by 65% and 69%, respectively, for the 300 mg/kg, while ALP was reduced by 67%.

#### 3.5.2. The histopathological Effect of 80% methanolic extraction of *P. africana*

The histopathological studies also supported the protective properties of the extract of leaves of *P. africana*. The areas of necrosis and ballooning degeneration of hepatocytes were observed in the CCl4-treated (toxic) group. The group of animals treated with *P. africana* showed a marked protective effect with decreased necrotic zones and hepatocellular degeneration. The photomicrographs of the liver sections are given in Fig 4.

**Figure 4:**
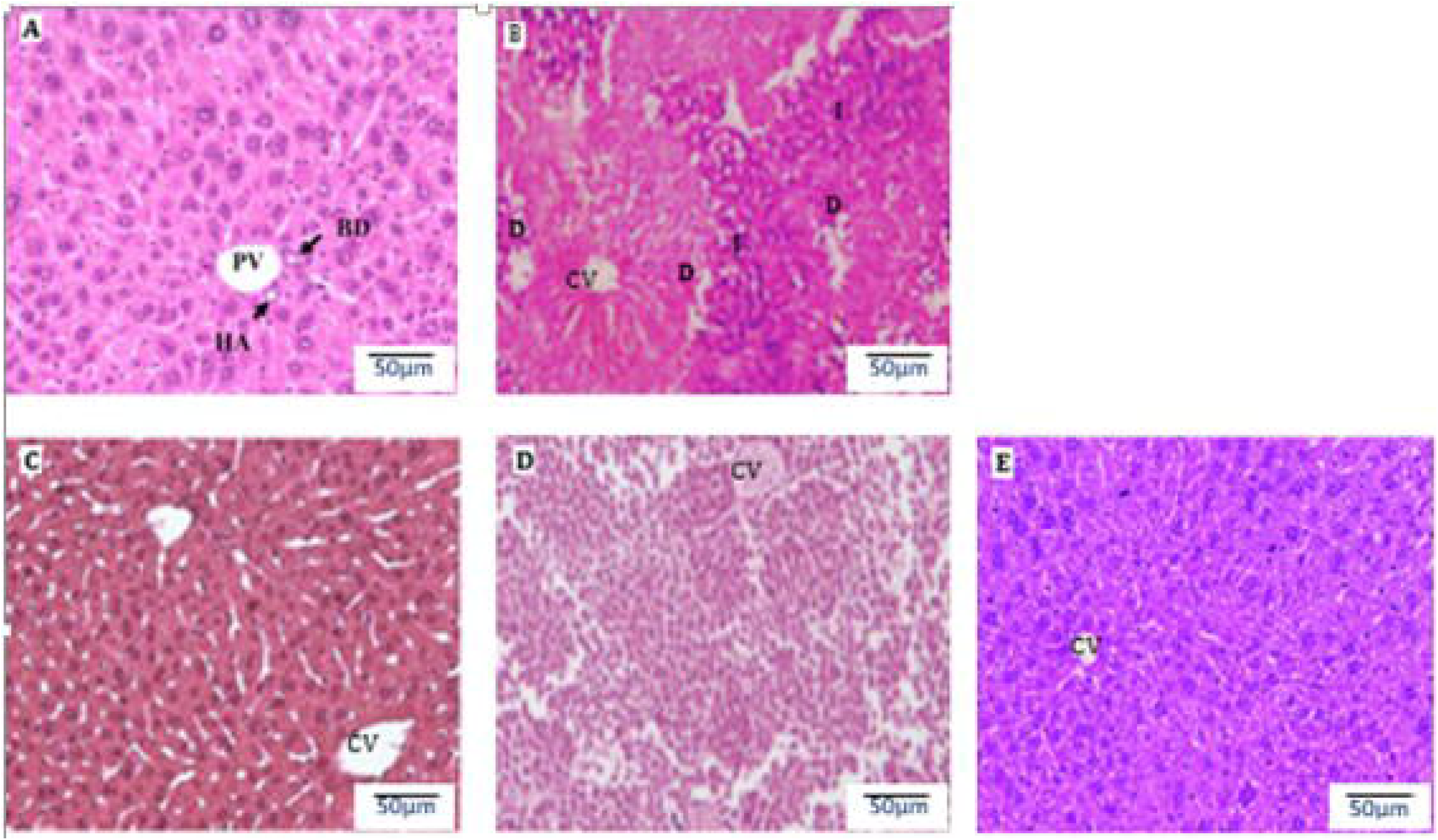
Paraffin sections of liver stained by hematoxylin and eosin for histopathological changes. (A) The liver section of the control group shows normal architecture. (B) Toxic liver, after treatment of CCl4 showing hepatic cell necrosis, fatty changes, or inflammatory cell infiltration. (C) The liver section was treated with CCl4+silymarin (100 mg/kg) preserving almost the normal structure of the hepatocytes. (D) The liver section was treated with CCl4 and P. africana (300 mg/kg) showing mild hepatic cell necrosis. (E) Liver section treated with CCl4 and P. africana (600 mg/kg) showing liver restoring to normal hepatic architecture. (X400).

## 4. Discussion

The hepatoprotective effect, toxicity test, and preliminary qualitative phytochemical tests of the crude extract made from *P. africana* leaves using 80% methanol were taken into consideration in this work. Consequently, the presence of many secondary metabolites in the crude extract of this plant species was demonstrated by the qualitative phytochemical assays. All of the crude extract taken from this plant showed good results for the presence of flavonoids, saponins, terpenoids, alkaloids, and tannins among the secondary metabolites seen in preliminary phytochemical screening, but negative results for anthraquinone glycoside.

This plant’s tannin content may indicate that it has antiparasitic, anticarcinogenic, and antimutagenic properties in addition to an astringent effect (26), reduction of HIV replication (27), and other properties (28). It has been demonstrated that terpenoids prevent the growth of certain microorganisms, including Candida albicans (14). An antioxidant impact linked to inflammation, autoimmune illnesses, cataracts, cancer, Parkinson’s disease, aging, and arteriosclerosis has been demonstrated by the extract’s flavonoid content (29). However, it has been discovered that the extract’s alkaloids have analgesic, antispasmodic, antihypertensive, anti-malarial, anticancer, and anti-inflammatory properties (21).

In Cameroon, a related earlier investigation *on P. africana* found the presence of tannins, terpenoids, steroids, and phenol (30). The stem bark extract of *P. africana* from Kenya’s woodland region contained alkaloids, glycosides, saponins, flavonoids, terpenoids, tannins, and phenolic compounds (21). The differences between *P. africana’s* chemical compositions in this study and those from earlier investigations might be due to variations in the geographical location of the plant, seasonal or chemo-type variations, and the type of particular phytochemical screening tests.

Toxicity studies or tests are undertaken to characterize any toxicity of a substance thereby ensuring safety (20). In all the cases, the toxic effects are usually manifested either in an acute, sub-acute, or chronic manner, and occur mostly as a result of an acute, sub-acute, or chronic exposure to toxic compounds by oral ingestion, inhalation, or absorption (21). In this study, the focus was on acute and sub-acute toxicity tests. The primary concern was to determine how the methanolic extracts of *P. africana* might be toxic after acute and sub-acute administration to Mice.

An acute toxicity test is a test that is conducted in a suitable animal species with a single dose and may be done for essentially all chemicals that are of any biological interest. Its purpose is to determine the symptomatology consequent to the administration of the compound and to determine the order of lethality of the compound (13). Accordingly, in the present study, the methanolic extract of *P. africana* resulted in one death after the third day of administration of the extract at a dose of 5000mg/kg. On the other hand symptoms like salivation, diarrhea, lethargy, and irritability were also observed following the administration of the extract on the remaining Mice at a dose of 5000mg/kg. Whereas such symptoms were subsided gradually and completely resolved after the first week of administration. A study of methanolic and aqueous extracts of *Prunus armeniaca* L which is similar in genius to *P*. *africana* did not show any signs and symptoms of toxicity and mortality up to 2000 mg/kg dose (22) and the researchers didn’t go further above the dose of 2000mg/kg.

The requirement to safeguard animal welfare was not taken into consideration when administering doses to animals over 5000 mg/kg. Animal testing at and above the (5000 mg/kg) range is discouraged and should only be taken into consideration in cases where there is a high probability that the results would directly benefit the health of humans or animals. No additional testing at higher dose levels should be carried out. (23). Therefore, based on the current data, it is anticipated that the oral LD50 of this plant’s methanolic leaf extract will be higher than 5000 mg/kg. Therefore, when taken orally at a dose lower than this, the plant’s extract might be regarded as harmless.

The weight growth of mice given dosages up to 5000 mg/kg throughout the 14-day acute toxicity investigation did not differ substantially (p>0.05) from the control groups. Furthermore, no gross pathological changes such as color, organ swelling, texture, and atrophy or hypertrophy were observed after a single administration of the extract as compared with the control group. Therefore, the overall weight gain in both experimental and control Mice and the absence of gross pathological changes indicated the good health status of the animals. In all groups, these results of the acute toxicity study go in line with another study of similar genius by Lenchyk LV. *et al*., (24) in which the body weights and gross anatomy of some of the internal organs including the liver and kidney of Mouse and Rats studied with acute administration of *Prunus domestica* leaves also did not significantly change as compared to the control.

There was no noticeable deviation in the behavior of the Mice administered with the low dose (600 mg/kg) as compared to that of the control group, and essentially all the experimental Mice remained healthy during the 28 weeks of sub-acute oral administration of *P. africana*. Moreover, no deaths were detected at both doses of this plant extract. Significant Changes in body weight have been used as an indicator of adverse effects of drugs and chemicals (25). Concerning this study, there was no significant change (p>0.05) in the weight of both male and female Mice throughout the study period, but there was a progressive non-significant weight increment in both experimental and the control groups. Non-significant Increment in body weight determines the positive health status of the animals (26). Therefore, the overall weight gain in both experimental and control Mice indicates a good health status of the experimental animals. Suggesting the lack of toxicity in all groups, these results of the sub-acute toxicity study go in line with another previous study by Dabe NE. *et al.* (27) in which the body weights of rats studied with acute and Sub-chronic oral administration of *A. afra* were not significantly changed as compared to the control.

Lu (25), states that as organ weight is impacted by the suppression of body weight, a significant shift in the relative organ weights of experimental and control animals is suggestive of toxicity. But in this investigation, there was no discernible shift in the weights of the liver or kidney, and a visual gross inspection revealed normal architecture, no color alterations, and no morphological abnormalities in the organs of either the experimental or control mice. This indicates that the sub-acute oral doses of *P. africana* extract administered had no effects on the organs of the Mice and was well tolerated over the 1-month study period.

Biochemical parameters were evaluated to obtain further toxicity-related information not detected by direct examination of organs and body weight analysis. Studies on biochemical parameters can easily reveal abnormalities in body metabolic processes. The blood profile usually provides important information on the response of the body to injury or lesion, deprivation, and stress (28).

Liver and renal function tests have a significant importance in evaluating changes produced by a toxicant. This is because of their response to clinical signs and systemic symptoms. To assess the possible toxic effect of a drug, evaluation of hepatic and renal function is primarily preferred as these organs are functionally predisposed. Elevated serum levels of enzymes produced by the liver or nitrogenous wastes to be excreted by the kidney might be an indication of their spillage into the bloodstream as a result of necrosis of the tissues (29). It was thus important to investigate the effect of *P. africana* on the function of these organs.

Liver chemistry tests include several serum chemistries that reflect liver function. The most commonly used serum liver chemistry tests include serum transaminases: ALT, AST, ALP, Gamma-glutamyl transpeptidase (GGT), bilirubin, and albumin. The major intracellular enzymes of the liver which are ALT and AST are useful as biomarkers for predicting possible toxicity (30). Any damage to the parenchymal liver cells will result in elevations in both of these transaminases (31). However, AST is not specific for the liver, because it is also present in other tissues like kidneys, heart, pancreas, skeletal muscle, brain, and red blood cells (3). The level of AST obtained in this study showed dose-dependent non-significant (p> 0.05) changes in the both male and female groups administered with *P. africana* extract at doses 600 & 1800mg/kg. These suggest that the Methanolic leaf extract of *P. africana* does not adversely affect the cell mitochondria as well as the other organelles in the cytoplasm; it may even stabilize these organelles (e.g. bring the level back to the day 1 levels).

Unlike AST, ALT is purely cytosolic and is more specific for hepatocytes. Serum transaminases, especially ALT, are the most important markers of hepatic injury(32). In the present study, there were no remarkable changes in ALT in male experimental mouse groups at doses of 600mg/kg and 1800mg/kg and there were non-significant increments of ALT at dose 1800mg/kg of all female groups administered with *P. africana.* These suggest that *P. africana* did not adversely affect the hepatocytes.

The liver is also the site of bilirubin synthesis. It is a catabolic end product from the breakdown of hem. The normal level is less than 1mg/dL (18 mmol/L)(33). In this study, there was no significant change of bilirubin on both experimental groups of male and female Mice. This indicated that the plant *P. africana* did not have a role in the derangement of liver function.

Urea & creatinine are the parameters to diagnose the functioning of the kidney. Changes in serum creatinine concentration more reliably reflect changes in Glomerular filtration rate (GFR) than do changes in serum urea concentrations (34). The rise in serum level of these chemicals indicates a decline or failure in renal function to filter waste products from the blood and excrete them in the urine (35). In the present study, the sub-acute oral administration of the extracts did not show significant alteration in the serum levels of urea and creatinine concentrations in Mice administered with both doses as compared to the controls. Thus, the absence of change in the levels of the above renal function markers suggests that the extract does not cause deterioration in renal function. This result is contrary to a study conducted by Chebaibi *et al.,* who investigated the toxicity of plant mixture used in the traditional treatment of edema and renal colic and showed a significant change in these parameters (36).

Histopathological examinations provide information to strengthen the results of other gross findings and biochemical parameters. Cell death due to toxicity (necrosis) or immunologically mediated occurs via apoptosis, in which isolated hepatocytes become shrunken, pyknotic, and intensely eosinophilic (37). Such hepatocyte damage did not occur in the present study as there was no focal necrosis, pyknosis, or enlarged or fragmented nucleus within the cytoplasm of hepatocytes at both 600 and 1800mg/kg doses.

On the other hand, there was mild mononuclear leukocytic cell infiltration near the central vein at a dose of 1800mg/kg of *P. africana* extract. However, such minor inflammatory changes obtained in this study were not accompanied by significant changes in any of the biochemical markers of the liver measured. The reason for the occurrence of the leukocytic infiltration is not clear, but it might be a response to parenchymal cell death with causes ranging from infectious agents, exposure to toxicants, generation of toxic metabolites, and tissue anoxia. In line with the present findings, a study conducted by Stephano HM *et al*. showed no significant change in general histological architectures of the liver parenchymal and non-parenchymal cells. Moreover, there were no effects on the levels of AST and ALT, which are considered to be sensitive indicators of hepatocellular damage and within limits that can provide a quantitative evaluation of the degree of damage to the liver (38). It is reasonable to deduce, therefore, that *P. africana* does not induce significant damage to the liver at both doses of the present study in both male and female Mice.

In line with the biochemical findings (the amount of creatinine and urea), the general histological architecture was not affected in any of the treatment groups as compared to the control, although minor lymphocytic infiltration was observed around the urinary pole in some kidney sections of Mice administered with the extraction only at 1800mg/kg body weight dose of *P. africana*. A similar study by Varsha R., *et al., on Prunus armeniaca* L. leaf extracts also showed the hepatic architecture was present in normal features of polygonal nucleus with nucleolus, abundant cytoplasm, and bilobed nucleus and showed no visible changes, disarrangement in hepatic cells (22).

CCl4 has been widely used in animal models to investigate chemical toxin-induced liver damage and as an excellent model to evaluate the efficacy of hepatoprotectants (39). The measure of a hepatoprotective medication’s protective properties is its capacity to lessen harmful effects or maintain the physiological processes of the liver that have been disrupted by a hepatotoxin (51). Therefore, by developing infiltration, vacuolization, and inflammation in the liver, mice given CCl4 in this study had larger liver weights. As a result, the mice’s weights dropped. On the other hand, mice pre- and post-treated with 80% methanol extracts of *P. africana* showed no significant difference in body weight, and both absolute and relative liver weight of mice as compared to the normal and standard control groups. A study conducted by Janey A. *et al*. showed similar types of results after the treatment of the hepatoprotective effects of ethanolic extraction of *Aquilaria agallocha* leaves against paracetamol-induced hepatotoxicity in rats (40).

Membrane damage results in the release of both cytosolic and endoplasmic enzymes, which show the presence of damage to liver structure and function (41). These are manifested as elevation in the levels of AST, ALT, and ALP (42). So, measuring the levels of these biomarkers of liver damage can reveal the hepatoprotective activity of the plant extract (43). In the present study, the 80% methanol extracts of *P. africana* showed a reduction in the levels of AST, ALT, and ALP in a dose-dependent manner. The 80% methanol extracts did produce a visible effect in all biomarkers of hepatic injury in its lower dose, but the higher doses were able to produce a more significant reduction in the levels of AST, ALT, and ALP. This could probably suggest that the lower dose might be below the minimum effective dose, which cannot elicit a significant reduction in liver enzyme levels and the other the higher dose might be large enough to cause a significant reduction. The percent reduction of biomarkers of liver injury showed that 600mg/kg of the Methanolic extracts exerted a nearly similar effect as that of the standard drug (silymarin). Pre- and post-treatment with 80% methanol extract at the dose of 600mg/kg largely modulated the severity of CCl4. In line with this, another Hepatoprotective study of the plant *Eclipta alba* against carbon tetrachloride-induced hepatotoxicity in albino rats showed that in the CCl4 administered Group: ALT, AST, ALP, total serum bilirubin were increased significantly whereas, administration of aqueous extract of *E. alba* at a dose of 250 mg/kg body weight showed statistically significant improvement in the serum ALT, AST, ALP, total serum bilirubin levels (44)

In this study, the levels of total bilirubin were used to assess liver synthetic and detoxification capability. A high concentration of bilirubin in serum is an indication of an increased erythrocyte degeneration rate (45). Due to the liver injury caused by the hepatotoxin, there is a defective excretion of bile by the liver which is reflected in their increased levels in serum (46). The oral administration of the methanolic leaves extraction of *P. africana* at 300 and 600mg/kg effectively reduced the serum total bilirubin levels which were comparable with both the normal and standard control groups.

Reinforcing the above-stated mechanisms, the histopathological study also supported the biochemical evidence for the hepatoprotection shown by *P. africana* in the same group. In the current investigation, the level of several biochemical measures considerably rose in the CCl4 toxic control groups, indicating the extent of hepatic injury as determined by histological evaluation. Methanolic extracts at dose levels of 300 and 600 mg/kg demonstrated a protective effect equivalent to that of the conventional medication Silymarine, which is 100 mg/kg. According to our findings, the methanolic extract of *P. africana* leaves may have hepatoprotective effects on mice’s liver damage induced by CCl4 through the following mechanisms: (1) inhibition of cytochrome P450-dependent oxygenase activity; (2) prevention of lipid peroxidation; and (3) stabilization of the hepatocyte membrane. After the phytochemical study, the plant extracts contain flavonoids, terpenoids, tannins, and steroids. The presence of flavanoids in all of our plant extracts may be responsible for its hepatoprotective activity (19,47–49).

## 5. Conclusion and Recommendations

In conclusion, the results of this study demonstrate that the extract of *P. africana* was effective for the prevention of CCl4-induced hepatic damage in Mice. Our results show that the hepatoprotective effects of *P. africana* extract may be due to marked changes in various parameters of liver enzyme, some physical parameters which proved that it’s a potent hepatoprotective in nature against the CCl4-induced liver toxicity. The present study thus justifies that, the use of *P. africana* in the treatment of liver diseases and points out that this plant warrants further detailed investigation as a promising hepatoprotective agent. However, the exact mechanism(s) and the active compound(s) involved in these effects need to be clarified in future studies. In addition, the observed results also raised the following concerns: First, although the sub-acute oral doses of *P. africana* extract used in this study did not produce any significant adverse effects in Mice, further studies using other organs like GIT, parts of the brain and CVS, and other animals are needed to be investigated. Second, the effect of various factors on the state of the plant material, such as the growth stage and maturity of the plant, the specific parts of the plant (leaves, roots, bark, flowers, seeds, etc), seasonal variation, and storage conditions may have effects on the chemical composition, and the toxicity studied may be extended in addressing these issues. Third, studies to determine the effects of *P. africana* on the fetus, in a pregnant animal, on the reproductive capacity of the animals, on the genetic system, and determine the ability of this plant to produce tumors (tumorigenicity and carcinogenicity tests) may also be investigated.

## Ethics approval

This study was carried out in strict accordance with the recommendations in the Guide for the Care and Use of Laboratory Animals of the National Institutes of Health. The protocol was approved by the Committee on the Ethics of Animal Experiments of the University of Mizan-Tepi (Protocol Number: MTU/CMCH/10034/2022). In accordance with the OECD 423 guideline, the study’s animals were also shielded from any needless agonizing or frightening circumstances (17). They were treated in accordance with the suggested protocol and given anesthesia for any uncomfortable procedures (18,50).

## Authors’ contributions

Each author contributed significantly and proportionately to the planning, execution of the study, gathering and analyzing the data. Each author contributed to the article’s drafting or critical revision for significant intellectual content. All authors committed to being accountable for every part of the work, giving their final consent for the version to be published, and submitting to this journal.

## Acknowledgment

We would first want to express our gratitude to the College of Health Sciences, the Offices of Research Directorate and Research Coordination, and the Vice President of Research and Community Services at Mizan-Tepi University for enabling us to participate in this study.

Second, we would like to extend our sincere gratitude to the traditional healers of Andracha Woreda, Sheka Zone, and Southwest Ethiopia for their gracious cooperation and courteous manner in giving the essential information and plant materials that were essential to the successful completion of the experiment Finally, we express our sincere gratitude to the Ethiopian Public Health Institute and Anatomy Department of Addis Ababa University for granting us study animals and access to their histology laboratory on schedule, respectively.

